# Oligogenic heterozygous inheritance of sperm abnormalities in mouse

**DOI:** 10.1101/2021.11.15.468601

**Authors:** Guillaume Martinez, Charles Coutton, Corinne Loeuillet, Caroline Cazin, Jana Muroňová, Magalie Boguenet, Emeline Lambert, Magali Dhellemmes, Geneviève Chevalier, Jean-Pascal Hograindleur, Charline Vilpreux, Yasmine Neirijnck, Zine-Eddine Kherraf, Jessica Escoffier, Serge Nef, Pierre F. Ray, Christophe Arnoult

## Abstract

Male infertility is an important health concern that is expected to have a major genetic etiology. Although high-throughput sequencing has linked gene defects to more than 50% of rare and severe sperm anomalies, less than 20% of common and moderate forms are explained. We hypothesized that this low success rate could at least be partly due to oligogenic defects – the accumulation of several rare heterozygous variants in distinct, but functionally connected, genes. Here, we compared fertility and sperm parameters in male mice harboring one to four heterozygous truncating mutations of genes linked to multiple morphological anomalies of the flagellum (MMAF) syndrome. Results indicated progressively deteriorating sperm morphology and motility with increasing numbers of heterozygous mutations. This first evidence of oligogenic inheritance in failed spermatogenesis strongly suggests that oligogenic heterozygosity could explain a significant proportion of asthenoteratozoospermia cases. The findings presented pave the way to further studies in mice and man.

## MAIN TEXT

Infertility is a major health concern affecting 15% of couples of reproductive-age worldwide^1, 2^. The infertility burden has increased globally for both genders in the past 30 years^3^, and now affects approximately 50 million couples worldwide^4^. Infertility is broadly treated by assisted reproductive technologies (ART), and today the number of individuals who were conceived by ART is close to 0.1% of the total world population, with over 8 million children already born following in vitro fertilization (IVF). Currently, ART is estimated to account for 1–6% of births in most countries^5^, with over 2.5 million cycles performed every year, resulting in over 500,000 births worldwide annually^6^. Despite this undeniable success, almost half the couples who seeks medical assistance for infertility fail to achieve successful pregnancy, and nearly 40% of infertile couples worldwide are simply diagnosed with unexplained or idiopathic infertility^7^. In the clinical context, few efforts are currently being made to understand and specifically address the underlying causes of a couple’s infertility because ART, as a palliative treatment, can often successfully rescue fertility even without a molecular diagnosis. Nevertheless, this absence of identification of the causes of infertility means that we lack alternative treatments for couples for whom current therapies are unsuccessful. Consequently, it is essential to improve the molecular diagnosis of infertility.

Human male infertility is a clinically heterogeneous condition with a complex etiology, in which genetic defects play a significant role. It is estimated that half of idiopathic cases of male infertility could be attributed to an as-yet unidentified genetic defect^8^. However, characterization of the molecular causes of male infertility represents a significant challenge, as over 4000 genes are thought to be involved in sperm production^9^. Over the past decade, significant progress has been made in gene identification thanks to the emergence of next- generation sequencing (NGS), and in functional gene validation thanks to new gene-editing techniques such as CRISPR/Cas9. NGS provides an inexpensive and rapid genetic approach through which to discover novel disease-associated genes^10^. It has proven to be a highly powerful tool in the research and diagnosis context of male infertility^11, 12^. In addition, validation of newly-identified variants through functional experiments has greatly benefited from the ability to generate mouse knock-out models using CRISPR technology^13^ and the use of novel model organisms like *Trypanosoma brucei* to study specific phenotypes, such as multiple morphological abnormalities of the [sperm] flagella (MMAF) syndrome^14–16^. These developments have resulted in a diagnostic yield, based on known genetic causes, explaining about 50% of cases of rare qualitative sperm defects like globozoospermia, acephalic, or MMAF syndromes^17–18^. In contrast, the diagnostic yield for quantitative sperm abnormalities such as oligozoospermia or azoospermia remains below 20%, even though these are the most common forms of male infertility^8, 12, 19^. To improve these low diagnostic yields, international consortia have been created to attempt to identify very low-frequency variants (https://gemini.conradlab.org/ and https://www.imigc.org/). In addition, several groups have started to assemble cohorts of patient-parent trios, aiming to identify de novo mutations causing male infertility as well as providing insight into dominant maternal inheritance^11, 20, 21^. Phenotypic heterogeneity and apparent incomplete penetrance were observed for some genetic alterations involved in male infertility^22–24^. These observations are difficult to reconcile with a model of Mendelian inheritance. We thus raised the possibility that the low diagnostic yield is partly due to the complex etiology of infertility, and hypothesized that some of the unsolved cases are due to oligogenic events, i.e., accumulation of several rare hypomorphic variants in distinct, functionally connected genes, and in particular to oligogenic heterozygosity. The molecular basis of oligogenicity is poorly understood. The main hypotheses are that two or more mutant proteins may act at different levels in the same intracellular pathway and could quantitatively contribute to its progressive dysfunction. When a critical threshold is reached, the disease phenotype would emerge. Alternatively, the mutant nonfunctional proteins produced may be part of the same multiprotein complex, and the presence of numerous pathogenic variants would increase the chance of the complex becoming compromised – leading to a progressive collapse of its cellular function^25^.

Here, we addressed the oligogenic heterozygosity hypothesis in male infertility using four specially-generated MMAF knock-out (KO) mouse models with autosomal recessive inheritance^14, 26^. Following extensive cross-breeding of our KO mouse lines, we produced lines harboring between one and four heterozygous truncating mutations. We assessed and compared fertility for these lines, analyzing both quantitative and qualitative sperm parameters, and performed a fine analysis of sperm nuclear morphology for all strains. Using this strategy, we were able to describe new genetic inheritance of sperm deficiencies.

## RESULTS

### Selection and characterization of individual MMAF mouse lines

To generate mice carrying up to four heterozygous truncating mutations, we first selected four lines carrying mutations in genes inducing a MMAF phenotype, namely *Cfap43*, *Cfap44*, *Armc2*, and *Ccdc146*. Three of these lines were already available and previously reported by our laboratory: a strain with a 4-bp deletion in exon 21 (delAAGG) for *Cfap43*^14^, a strain with a 7-bp insertion in exon 3 (InsTCAGATA) for *Cfap44*^14^, and a strain with a one-nucleotide duplication in exon 4 (DupT) for *Armc2*^26^, inducing a translational frameshift that leads to the production of a truncated protein. We generated the fourth strain using CRISPR-Cas9 technology as described in the material and methods section, inducing a 4-bp deletion in exon 2 (delTTCG) of the *Ccdc146* gene (Supplementary Figure 1). A study describing how this mutation in the *Ccdc146* gene affects spermatogenesis is currently under review elsewhere (for the reviewers only, we provide the complete phenotype for the Ccdc146 KO mouse strain, demonstrating its role in MMAF syndrome in mice).

We first confirmed the MMAF phenotypes for all four strains. Sperm from all homozygous KO male mice displayed more than 95% morphological abnormalities of the flagellum including coiled, bent, irregular, short or/and absent flagella (Figures 1-4). We then analyzed sperm morphology (head and flagellum) in heterozygous animals by optical microscopy. The four strains fell into two categories. For the two strains targeting *Cfap43* and *Cfap44*, *Cfap43^+/-^*, or *Cfap44^+/-^* heterozygous males had slightly higher rates of abnormalities than wild-type mice (Figure 1B and 2B; *Cfap43*: t=-2.79, df=6.13, p-value=0.03; *Cfap44*: t=-8.80, df=6.14, p-value=0.0001). These results were in accordance with previous observations^14^. In contrast, for strains harboring heterozygous mutations in *Armc2* and *Ccdc146*, no significant differences were observed with respect to wild-type males (Figure 3B and 4B).

**Figure 1.**
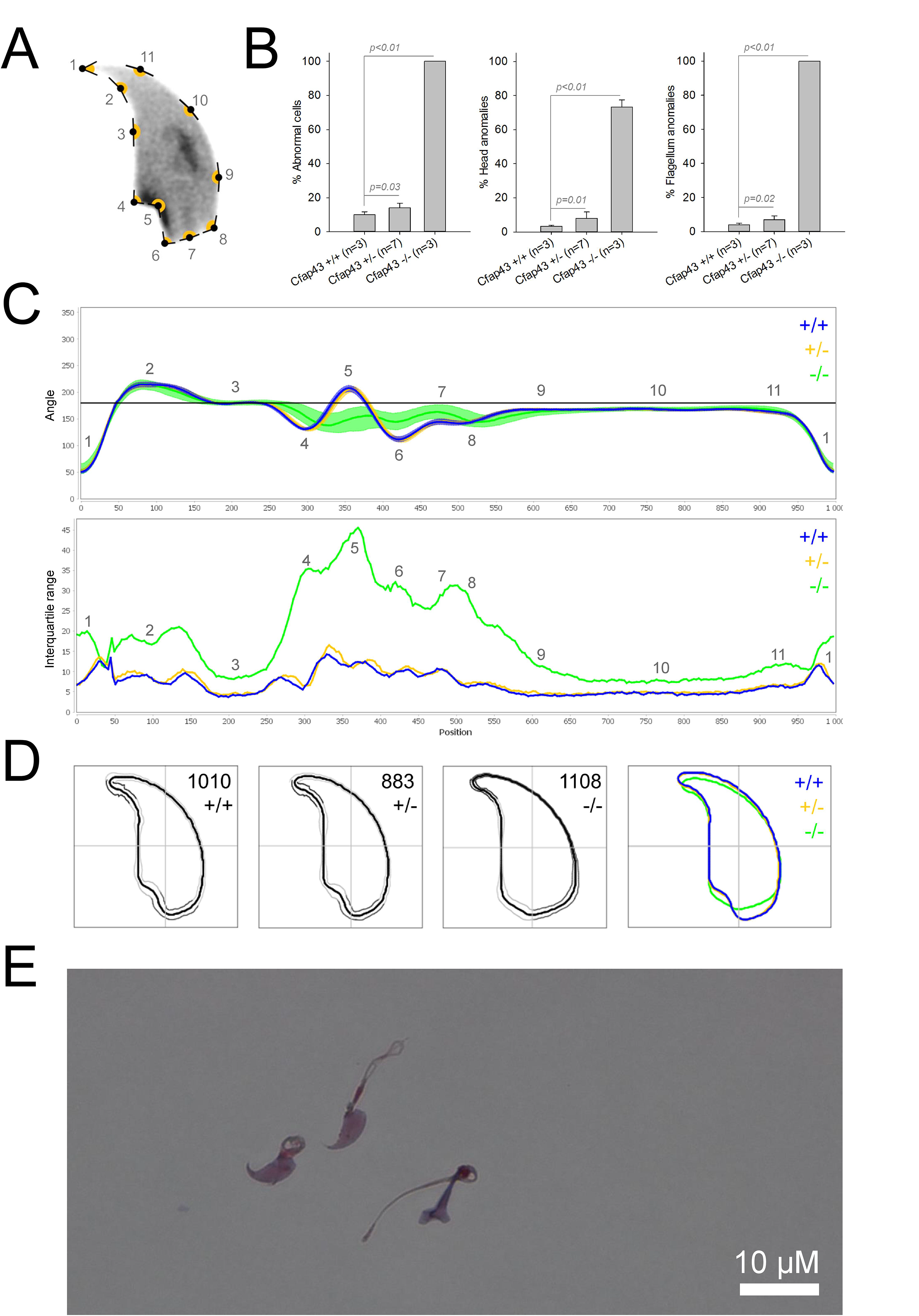
Sperm morphology analysis for the *Cfap43* mouse strain. **A.** Graphical representation of the location and numbering of the regions of interest presented in the lineage profiles (from ^27^) with: 1-tip; 2-under-hook concavity; 3-vertical; 4-ventral angle; 5-tail socket; 6-caudal bulge; 7-caudal base; 8-dorsal angle; 9-11-acrosomal curve. **B.** Histogram showing proportions of morphological anomalies (mean ± SD) for each *Cfap43* genotype. Statistical significance was assessed by applying an unpaired *t*-test; *p*-values are indicated. **C.** Angle profiles (top) and variability profiles (bottom) from *Cfap43^+/+^* (blue), *Cfap43^+/-^* (yellow), and *Cfap43^-/-^* (green) mice. The x axis represents an index of the percentage of the total perimeter as measured counterclockwise from the apex of the sperm hook. The y axis corresponding to the angle profile represents the interior angle measured across a sliding window centered on each index location. The y axis corresponding to the variability profile represents the Interquartile Range (IQR) value (the difference between the 75^th^ and 25^th^ percentiles). Specific regions of the nuclei are mapped on both axes as indicated in A. **D.** Consensus nuclear outlines for each genotype alongside a merged consensus nucleus (blue=*Cfap43^+/+^*, yellow= *Cfap43^+/-^*, green= *Cfap43^-/-^*). The numbers assigned to each consensus outline correspond to the number of nuclei processed per condition. **E.** Optical microscopy analysis showing a representative MMAF phenotype for *Cfap43* KO mice (scale bar, 10 µM).

**Figure 2.**
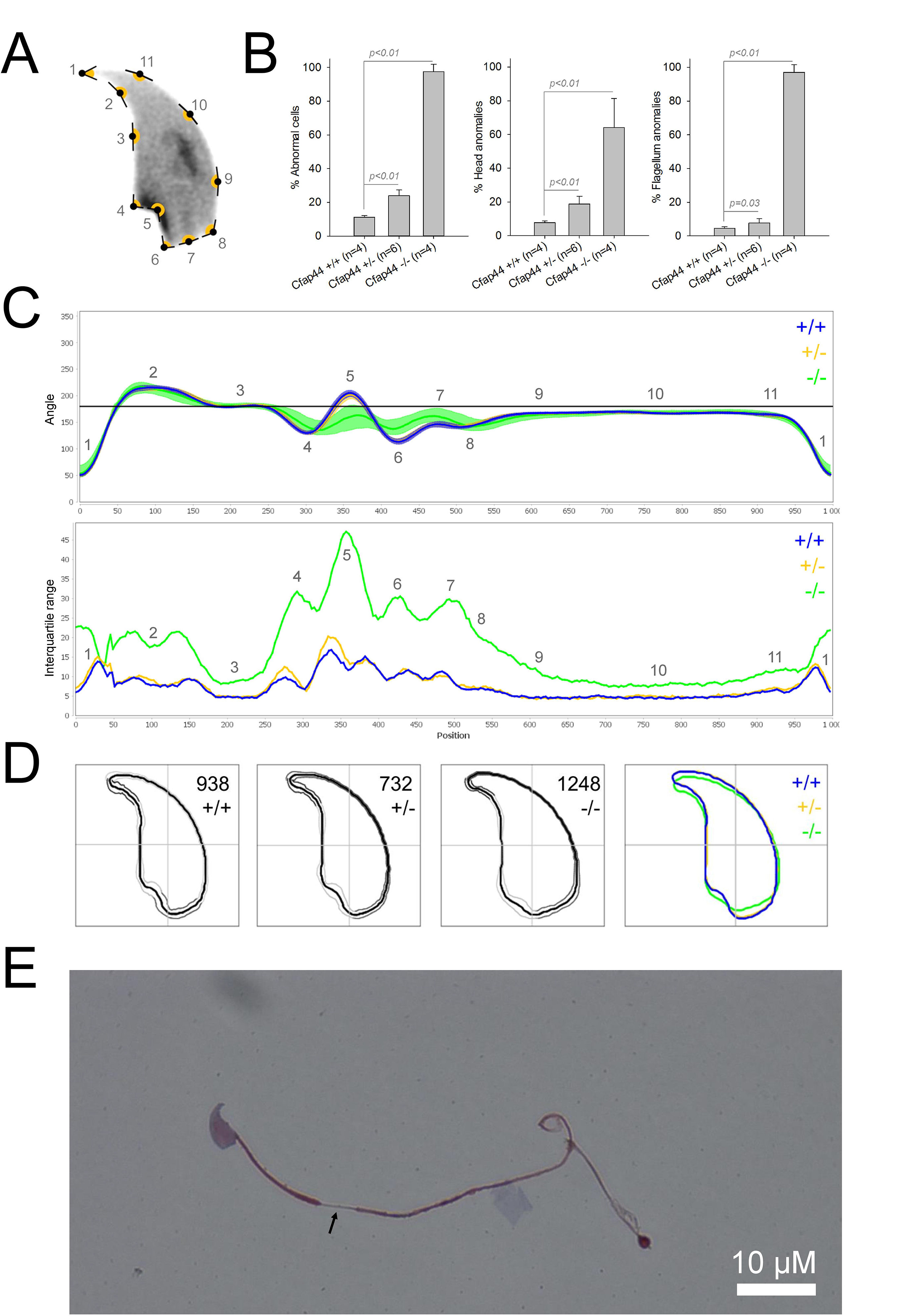
Sperm morphology analysis for the *Cfap44* mouse strain. **A.** Graphical representation of the location and numbering of the regions of interest presented in the lineage profiles (from ^27^) with: 1-tip; 2-under-hook concavity; 3-vertical; 4-ventral angle; 5-tail socket; 6-caudal bulge; 7-caudal base; 8-dorsal angle; 9-11-acrosomal curve. **B.** Histogram showing proportions of morphological anomalies (mean ± SD) for each *Cfap44* genotype. Statistical significance was assessed by applying an unpaired *t*-test; *p*-values are indicated. **C.** Angle profiles (top) and variability profiles (bottom) for *Cfap44^+/+^* (blue), *Cfap44^+/-^* (yellow), and *Cfap44^-/-^* (green) mice. The x axis represents an index of the percentage of the total perimeter, as measured counterclockwise from the apex of the sperm hook. The y axis corresponding to the angle profile is the interior angle measured across a sliding window centered on each index location. The y axis corresponding to the variability profile represents the Interquartile Range (IQR) value (the difference between the 75^th^ and 25^th^ percentiles). Specific regions of the nuclei are mapped on both axes as indicated in Fig. 1A. **D.** Consensus nuclear outlines for each genotype alongside a merged consensus nucleus (blue=*Cfap44^+/+^*, yellow= *Cfap44^+/-^*, green= *Cfap44^-/-^*). The numbers assigned to each consensus outline correspond to the number of nuclei processed per condition. **E.** Optical microscopy analysis showing a representative MMAF phenotype for *Cfap44* KO mice (scale bar, 10 µM).

**Figure 3.**
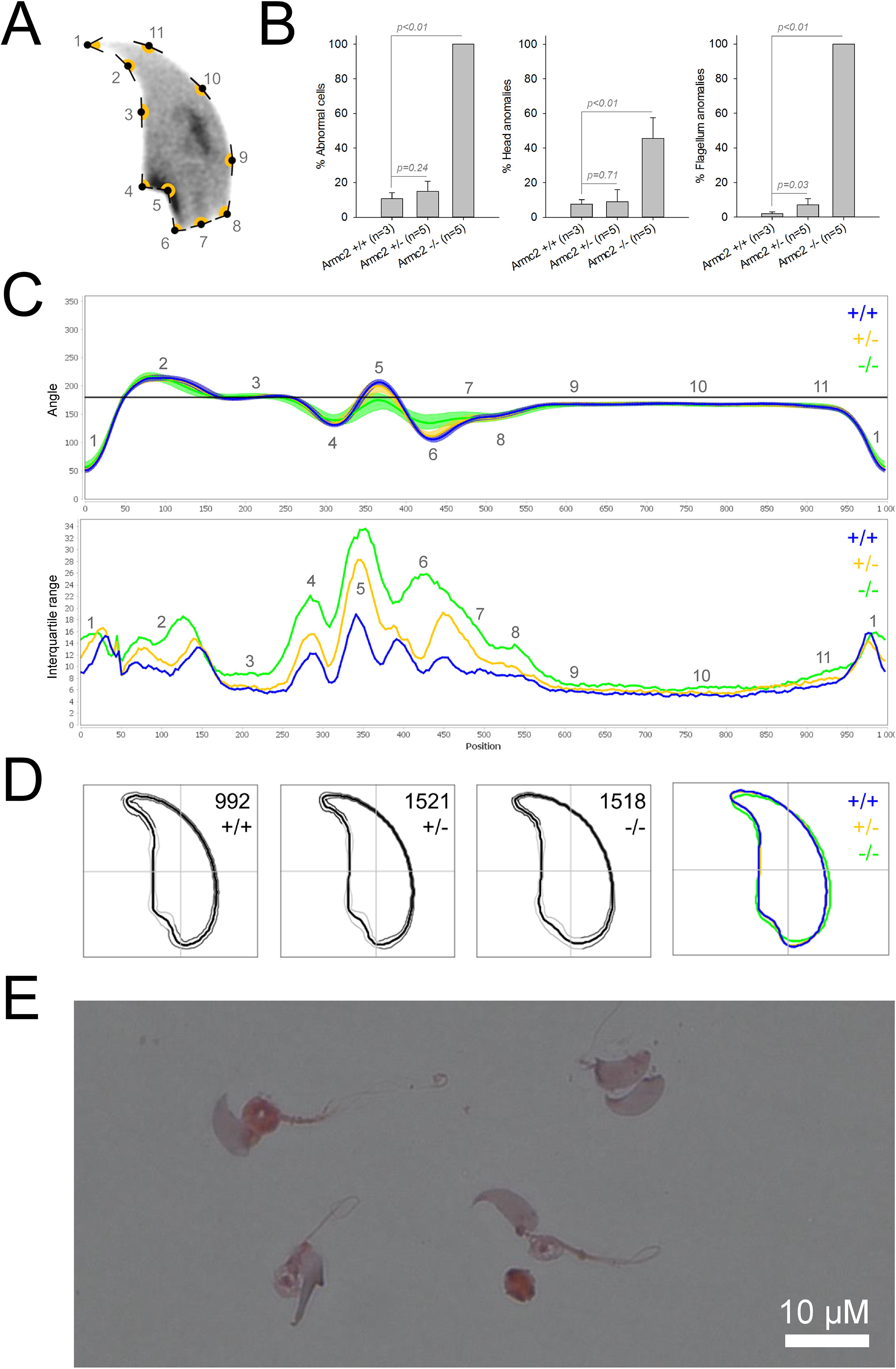
Sperm morphology analysis for the *Armc2* mouse strain. **A.** Graphical representation of the location and numbering of the regions of interest presented in the lineage profiles (from ^27^) with: 1-tip; 2-under-hook concavity; 3-vertical; 4-ventral angle; 5-tail socket; 6-caudal bulge; 7-caudal base; 8-dorsal angle; 9-11-acrosomal curve. **B.** Histogram showing proportions of morphological anomalies (mean ± SD) for each *Armc2* genotype. Statistical significance was assessed by applying an unpaired *t*-test; *p*-values are indicated. **C.** Angle profiles (top) and variability profiles (bottom) for *Armc2^+/+^* (blue), *Armc2^+/-^* (yellow), and *Armc2^-/-^* (green) mice. The x axis represents an index of the percentage of the total perimeter, as measured counterclockwise from the apex of the sperm hook. The y axis corresponding to the angle profile represents the interior angle measured across a sliding window centered on each index location. The y axis corresponding to the variability profile represents the Interquartile Range (IQR) value (the difference between the 75^th^ and 25^th^ percentiles). Specific regions of the nuclei are mapped on both axes as indicated in Fig. 1A. **D.** Consensus nuclear outlines for each genotype alongside a merged consensus nucleus (blue=*Armc2^+/+^*, yellow=*Armc2^+/-^*, green=*Armc2^-/-^*). The numbers assigned to each consensus outline correspond to the number of nuclei processed per condition. **E.** Optical microscopy analysis showing a representative MMAF phenotype for *Cfap44* KO mice (scale bar, 10 µM).

**Figure 4.**
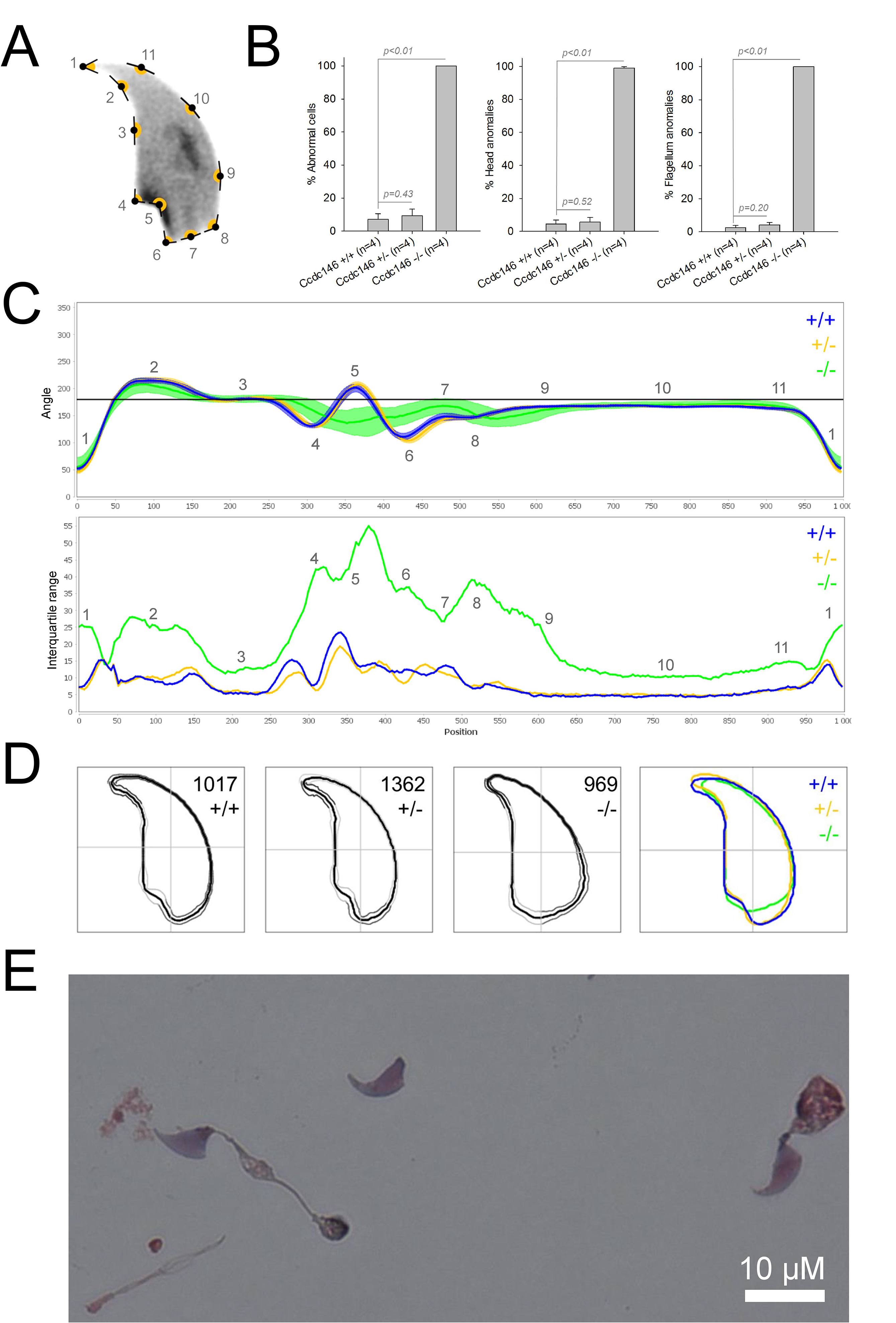
Sperm morphology analysis for the *Ccdc146* mouse strain. **A.** Graphical representation of the location and numbering of the regions of interest presented in the lineage profiles (from ^27^) with: 1-tip; 2-under-hook concavity; 3-vertical; 4-ventral angle; 5-tail socket; 6-caudal bulge; 7-caudal base; 8-dorsal angle; 9-11-acrosomal curve. **B.** Histogram showing proportions of morphological anomalies (mean ± SD) for each *Ccdc146* genotype. Statistical significance was assessed by applying an unpaired *t*-test; *p*-values are indicated. **C.** Angle profiles (top) and variability profiles (bottom) for *Ccdc146^+/+^* (blue), *Ccdc146^+/-^* (yellow) and *Ccdc146^-/-^* (green) mice. The x axis represents an index of the percentage of the total perimeter as measured counterclockwise from the apex of the sperm hook. The y axis corresponding to the angle profile is the interior angle measured across a sliding window centered on each index location. The y axis corresponding to the variability profile represents the Interquartile Range (IQR) (the difference between the 75^th^ and 25^th^ percentiles). Specific regions of the nuclei are mapped on both axes as indicated in Fig. 1A. **D.** Consensus nuclear outlines for each genotype alongside a merged consensus nucleus (blue=*Ccdc146^+/+^*, yellow=*Ccdc146^+/-^*, green=*Ccdc146^-/-^*). The numbers assigned to each consensus outline correspond to the number of nuclei processed per condition. **E.** Optical microscopy analysis showing a typical MMAF phenotype for *Ccdc146* KO mice (scale bar, 10 µM).

Because head morphology defects may be subtle and difficult to detect by visual observation, we applied a newly-developed method, involving the use of Nuclear Morphology Analysis Software (NMAS)^27^. The method is described in detail in the material and methods section. The nuclear morphologies of sperm from each genotype (WT, heterozygotes, and homozygotes) were characterized. For all corresponding wild-type strains, shape modeling gave extremely similar consensus and angle profiles for nuclei, highlighting the common genetic background of the KO animals (Supplementary figure 2). For heterozygous *Cfap43^+/-^*, *Cfap44^+/-^*, and *Ccdc146^+/-^* individuals, the angle profiles were very similar to WT profiles (Figure 1C, 2C, and 4C). In contrast, *Armc2^+/-^* individuals showed a slightly modified angle profile compared to the WT profile, due to a narrower caudal base inducing a more pronounced caudal bulge and a reduced dorsal angle (Figure 3C). When the profiles for all heterozygous males were compared, a very similar nuclear morphology with nearly identical angle and variability profiles was found, apart from for *Armc2^+/-^*, which had slightly more variability than the other lines despite a similar angle profile (Supplementary Figure 3).

We also compared these profiles with those of the corresponding KO mice, which displayed very unique and specific patterns (Figure 1C, 2C, 3C, 4C). Briefly, *Cfap43^-/-^* and *Cfap44^-/-^* angle profiles both displayed a flattening from the ventral angle to the dorsal angle leading to their characteristic “pepper” shape. The lines for both profiles merged over the majority of regions such as the under-hook concavity and the acrosomal curve, and showed intermediate impairment, greater than that observed in the *Armc2^-/-^* line and less than that observed in the *Ccdc146^-/-^* line. The angle profile for *Armc2^-/-^* individuals was similar to that of wild-type individuals at the tip, dorsal angle, and acrosomal curve, as these regions do not appear to be affected by the mutation. Under-hook concavity, ventral angle, and tail socket regions were slightly affected, although less than in other lineages. In contrast to the other lineages, the caudal base was markedly shortened, with a significant flattening observed. For the *Ccdc146^-/-^* line, with the exception of the ventral-vertical region, which was unaffected in any lineage, all regions of the angle profile were strongly impacted. The dorsal angle was completely absorbed into the acrosomal curve, and the under-hook concavity, ventral angle and caudal bulge regions were flattened. The tail socket region even displayed an inverted angular profile compared to the other lineages. Overall, the profile of *Ccdc146^-/-^* showed considerable variability, and complete remodeling of the angle profile with, among other changes, total inversion of the curve at position 330-400, corresponding to complete disappearance of the tail socket. Comparison of the profiles for KO sperm (Supplementary Figure 4) showed that *Cfap43^-/-^* and *Cfap44^-/-^* individuals displayed similar alterations (general enlargement of the head, rounding of the base at the expense of the flagellum insertion, etc.) resulting in close consensus; the other two lines were quite distinct. Whereas it was the most affected heterozygous line, *Armc2^-/-^* mice displayed the mildest alterations to nuclear morphology and variability of the four lines. KO animals nonetheless retained specific patterns, including modification of the hook curve and a specific slope of the base profile. Finally, *Ccdc146^-/-^* individuals displayed extremely severe alterations, with almost all of their nuclei presenting a triangular shape (Figure 4DE). These results demonstrate for the first time that absence of MMAF genes not only affects sperm flagellum biogenesis but also has a dramatic impact on sperm head morphology.

From these extensively characterized lines, we then proceeded to generate individuals harboring multiple mutations.

### Impact of accumulation of heterozygous mutations

All individuals harboring between one and four heterozygous truncating mutations were generated by standard cross-breeding of the four lines. A considerable number of generations were produced over several years. Due to time and financial constraints, only one combination of multiple heterozygous lines was created and analyzed. Double heterozygotes were obtained by crossing *Cfap43* and *Cfap44* KO mouse lines. Triple heterozygous animals were mutated for *Cfap43*, *Cfap44*, and *Armc2*; and the quadruple heterozygous line also had the *Ccdc146* mutation (Figure 5).

**Figure 5.**
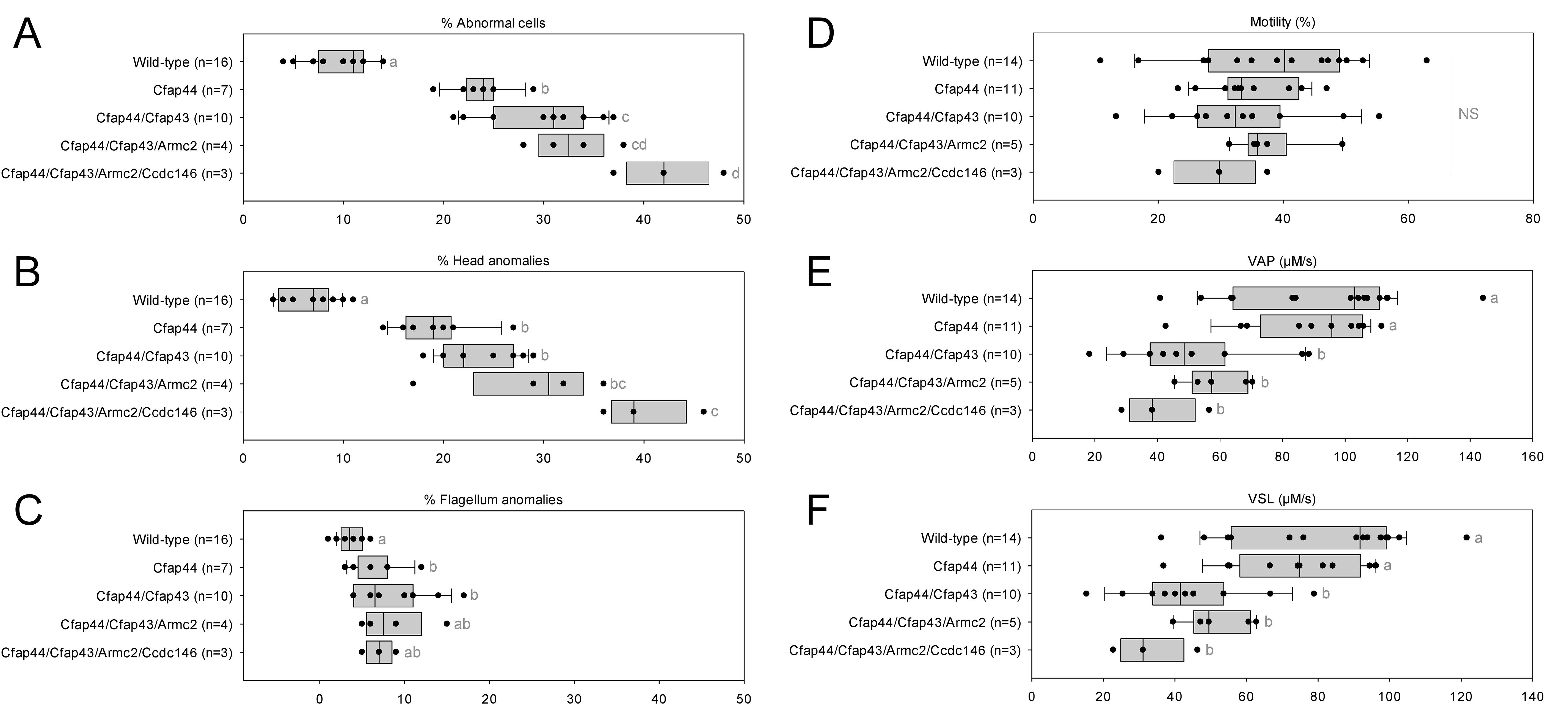
Increasing the number of heterozygous mutated genes involved in the MMAF syndrome has a drastic effect on sperm morphology and motility. (A) Sperm morphological defects, (B) head anomalies, (C) flagellum defects, showing only slight impact. Increasing the number of heterozygous gene mutations had little effect on (D) Percentage of motile sperm, but considerably reduced sperm motility parameters including (E) average path velocity (VAP), and (F) curvilinear velocity (VCL). All data are presented simultaneously as box-plot and individual datapoints. Statistical significance was assessed using an unpaired *t*- test, and significantly different populations are identified by distinct letters. The corresponding statistical data can be found in Supplementary Tables 1-2.

The accumulation of heterozygous mutations on the four selected genes involved in MMAF syndromes induced qualitative spermatogenesis defects, with increased numbers of morphological anomalies (Figure 5A), in particular defects of the head (Figure 5B). In contrast, flagellar anomalies were not amplified as the number of mutated genes increased (Figure 5C). Nevertheless, accumulation of mutations had a negative impact on sperm motility parameters. Although the overall percentage of motile cells decreased only slowly with increasing numbers of mutations (Figure 5D), the quality of sperm movement was strongly affected (Figure 5E-5F). Thus, for sperm bearing two mutations, the average sperm velocity and straight-line velocity were halved compared to control individuals, and the decreasing trend continued as mutations accumulated. It is worth noting that adding *Ccdc146* and *Armc2* heterozygous mutations aggravated the phenotype, even though the heterozygous mutations alone had no impact on the sperm phenotype observed by optical microscopy (Figure 3B and 4B).

From the crosses performed to generate multi-heterozygous animals, other combinations of heterozygous mutations were obtained. The sperm parameters of the corresponding animals were also phenotyped. Interestingly, rates of morphological defect (Figure 6A-C, red dots) and motility parameters (Figure 6D-F, red dots) were very similar whatever the gene combinations. Taken together, these results show that it is not the specific combination of heterozygous mutations that leads to altered sperm morphology and sperm motility parameters, but rather their accumulation.

**Figure 6.**
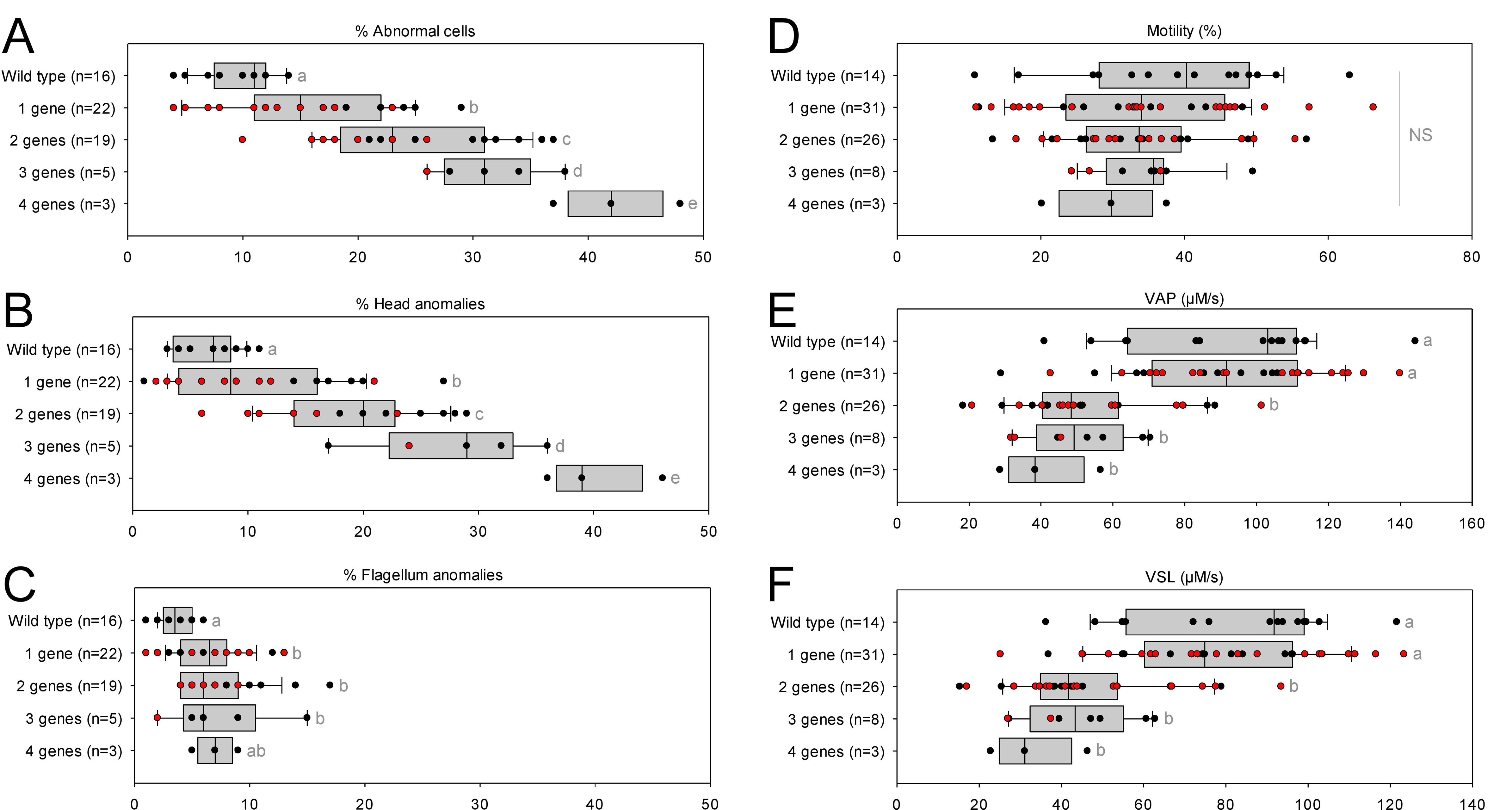
Very similar head morphology defects and decreased motility parameters, whatever the combination of mutated genes. Black dots correspond to the different combinations (2 = Cfap43 and Cfap44; 3 = Cfap43, Cfap44 and Armc2; 4 = Cfap43, Cfap44 and Armc2 and Ccdc146) of mutations presented in figure 5; red dots correspond to alternative combinations obtained with the same four genes. The range of (A) sperm morphological defects, (B) head anomalies, and (C) flagellum defects was similar for black and red dots. Likewise, (D) percentage of motile sperm, (E) average path velocity (VAP) and (F) curvilinear velocity (VCL), showed similar ranges for black and red dots. All data are presented simultaneously as box-plot and individual datapoints. Statistical significance of differences between means calculated for all black and red dots were assessed by applying an unpaired *t*-test. Statistically different populations are identified by distinct letters. The corresponding statistical data can be found in Supplementary Tables 3-4.

Despite a marked alteration of spermatocytograms, accumulation of mutations did not have a significant effect on quantitative spermatogenesis defects. For example, sperm production – represented by testis weight and sperm concentration – and overall fertility of the animals – based on the number of pups per litter and interval between litters – were not affected, whatever the combination and number of mutations (Figure 7).

**Figure 7.**
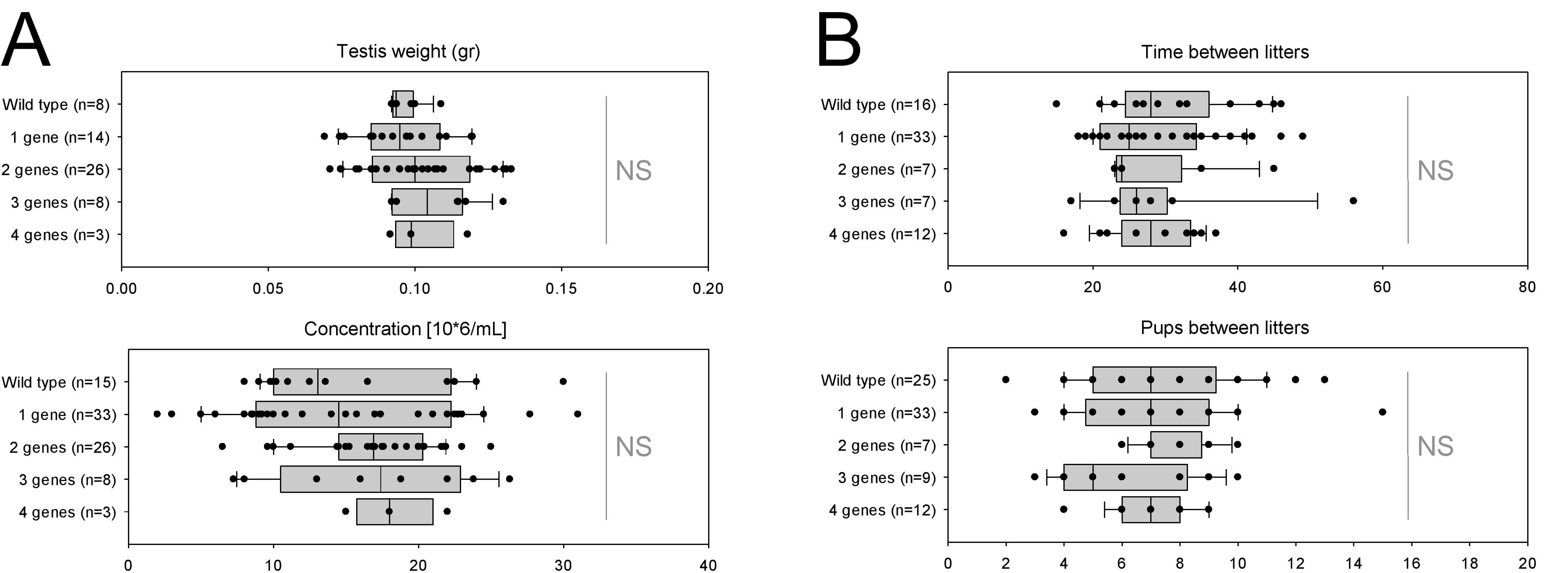
Increasing the number of mutated genes has little effect on overall fertility. (A) sperm production data (B) overall fertility of animals (measured as the interval between two litters and the number of pups per litter). All data are presented simultaneously as box-plot and individual datapoints. Statistical significance was assessed using an unpaired *t*-test, and significantly different populations are identified by distinct letters. The corresponding statistical data can be found in Supplementary Tables 5-6.

As for mouse lines bearing single mutations, we then used NMAS to help characterize head anomalies in multi-heterozygote mutated lines. Our results indicated a striking progressive deterioration in sperm head morphology as mutations accumulated (Figure 8). Each additional mutation progressively and significantly increased the variability and severity of the morphological defects observed in the nucleus, negatively influencing angle and consensus profiles. This negative effect was cumulative, and if a threshold or plateau effect exists, it was not reached upon accumulation of four mutations.

**Figure 8.**
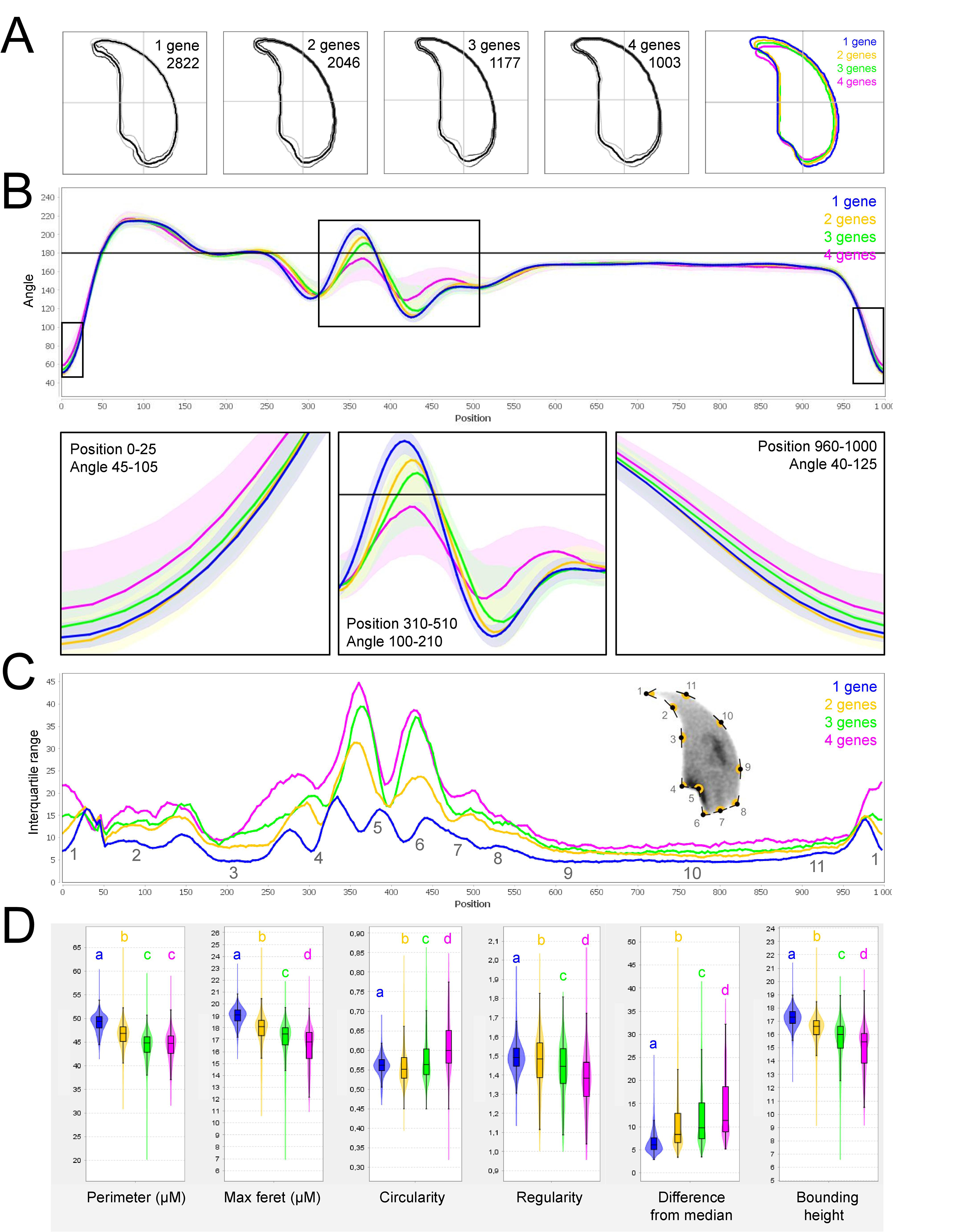
Fine nuclear morphology analysis of multi-mutant mice with one to four heterozygous mutations. **A.** Consensus nuclear outlines for each strain alongside a merged consensus nucleus (blue=one mutation, yellow=two mutations, green=three mutations and pink=four mutations). The numbers assigned to each consensus outline correspond to the number of nuclei processed per condition. **B.** Angle profiles for each strain focusing (black boxes) on positions of specific interest. The x axis represents an index of the percentage of the total perimeter, as measured counterclockwise from the apex of the sperm hook. The y axis shows the interior angle measured across a sliding window centered on each index location. **C.** Variability profiles for each strain. The x axis is the same as for the angle profile, and the y axis represents the Interquartile Range (IQR) (the difference between the 75^th^ and 25^th^ percentiles). Specific regions of the nuclei are mapped on the profile and the graphical representation (from ^27^), with: 1-tip; 2-under-hook concavity; 3-vertical; 4-ventral angle; 5-tail socket; 6-caudal bulge; 7-caudal base; 8-dorsal angle; 9-11-acrosomal curve. **D.** Violin plots of nuclear parameters for each strain. Statistical significance of differences between populations were assessed by the software, applying a Mann-Whitney U test; significantly different populations are identified by distinct letters.

To extend our analysis, we then used the software to analyze sperm sub-populations by performing unbiased nuclear morphology categorization. Clustering based on angle profiles revealed a total of ten sub-groups of nuclei shape, which matched with the usual shapes of normal and abnormal mouse sperm, defined more than 30 years ago^28^. Mutation-accumulation progressively increased the frequency of all abnormal shapes and decreased the frequency of normal forms. The most unstructured forms were associated with the highest number of mutations (Supplementary figure 5).

## DISCUSSION

The aim of this study was to determine whether the accumulation of several rare heterozygous variants in functionally connected genes affected fertility and sperm parameters in mice. Our results clearly demonstrated that spermatogenesis failure can arise from oligogenic heterozygosity in mice. Males bearing increased numbers of heterozygote mutations in genes involved in MMAF syndrome exhibited altered spermatocytogram, with a greater proportion of abnormal sperm and decreased sperm motility. However, these alterations had no effect on male fertility, suggesting that we had not reached the threshold leading to the complete collapse of the male reproductive function. This concept of threshold is strongly associated with oligogenicity. It is defined by the number of mutations within the same multiprotein complex or intracellular pathway beyond which a disease phenotype will be observed^29^. In this study, we induced mutations in four genes coding for proteins participating in flagella formation and function. Although we did observe a negative cumulative burden, we did not reach the complex or system threshold that would lead to the emergence of a dichotomous severe infertility phenotype. Nevertheless, mice have particularly efficient sperm production and their male and female reproductive functions are generally robust. A similar mutational burden in humans - known to have the highest number of sperm defect among primates^30^ – could, however, be sufficient to reach the threshold for fertility collapse. It is worth noting that the probability of accumulating this type of heterozygous mutations in testis is considerable, because i) thousands of genes are necessary to achieve spermatogenesis^9^, ii) expression of most spermatogenesis-associated genes is restricted to or strongly enriched in the testis^31^, and consequently iii) the risk of life-threatening impact of mutations is limited.

Moreover, mutant gene products retaining some residual function could be influenced by additional systemic perturbation (see review in ^32^) that would lead to system collapse. For instance, environmental factors commonly responsible for milder alterations to spermatogenesis could play an important role by severely aggravating the genetic burden on the system. This type of multi-factorial input could explain the phenotypic continuum observed in patients with idiopathic infertility.

In this article, we showed that the haplo-insufficiency of several genes involved in flagellum biogenesis and associated with MMAF syndrome leads to head defects. The impact of the accumulation of mutations in the heterozygous state on the morphology of the head is in agreement with our results (presented above and previously published^14^) showing that the complete lack of these genes in KO males also strongly alters head patterning. Unexpectedly, accumulation of heterozygous mutations led to marked head defects without obvious morphological defects of the flagellum. This observation suggests that head shape is more prone to collapse than flagellum biogenesis. For the first time, we were able to objectively document these defects using NMAS. This original method allows objective comparison of the impact of mutations on sperm head morphology, and represents a breakthrough in sperm morphology assessment in a pathological context. The NMAS method was also used to analyze head morphology in sperm produced by KO males. We found that sperm from *Cfap43^-/-^* and *Cfap44^-/-^* mice predominantly displayed a “pepper” shape – characterized by a strong broadening of the base and a reduced size of the flagellum insertion notch. *Armc2^-/-^* sperm show moderate enlargement extending along the entire length of the head, including the hook, and an increase in circularity resulting in a more rounded appearance. *Ccdc146^-/-^* sperm presented a total disorganization of the base with complete erasure of the flagellum insertion notch, and an overall decrease in size, resulting in heads with a triangular aspect characteristic of the “claw” shape. The reasons why mutations in flagellar proteins have such a strong impact on head morphology remain unclear, however the effect on sperm head shape was gene-dependent, with a milder effect observed for *Armc2* and a stronger effect observed for *Cfap43*, *Cfap44*, and *Ccd146*. This result was notable as the flagellum phenotype of *Armc2* is as severe as that observed with mutation of the other genes. Consequently, sperm head defects are not due to failed flagellum biogenesis. Rather, we hypothesize that some proteins contribute not only to flagellum biogenesis, but also to various cytoskeletal components including the manchette, a key organelle in sperm head shaping^33^. For instance, a recent report showed that lack of Cfap43 affects intra-manchette transport^34^. Thus, compared to the other proteins, Armc2 may be less involved in functions other than flagellum biogenesis. Therefore, we recommend that head morphology should not be overlooked when studying MMAF syndromes, despite an obvious focus on flagellum anomalies.

Among the proteins encoded by previously identified MMAF genes, some belong to complexes involved in intraflagellar transport^35–37^, protein degradation^38^ or unknown processes^26, 39^ that could affect head formation, and subsequently sperm DNA. As there is a strong relationship between cytoskeletal and chromosomal effects, the question of the potential impact of the accumulation of mutations on sperm chromatin organization arises. Another interesting question would be to investigate whether the damaged heads correspond to those carrying the mutated alleles. Future studies will be eagerly awaited to elucidate the molecular basis of oligogenicity in male infertility.

Moreover, as descriptions of animal reproductive phenotypes are becoming increasingly sophisticated and standardized^40^, emerging tools such as the software used in this study to analyze nuclear morphology should be used to extensively study reproductive phenotypes, including studies of MMAF syndromes, to accurately document head defects.

Despite a strong focus on sperm head defects, we also showed that sperm motility was altered in the presence of ≥2 heterozygote mutations. Thus, the function of the flagellum is affected, and an absence of morphological defects should therefore not be considered synonymous with absence of functional defects. More importantly, this result strongly suggests for the first time that idiopathic human asthenozoospermia may be due to an accumulation of heterozygous mutations in genes known to be involved in flagellum biogenesis.

To date, the inheritance pattern of isolated male infertility was only known to be Mendelian – i.e., based on a single locus – and the possibility of oligogenic inheritance had not been explored. In contrast, an oligogenic etiology for female infertility has been proposed to be associated with primary ovarian insufficiency (POI) by several groups, in particular due to the identification of heterozygous mutations in several genes associated with POI^41^. Oligogenic inheritance has also previously been suggested in another reproductive disorder: congenital hypogonadotropic hypogonadism^42^, as well as in several non-reproductive disorders^43^. Our results demonstrate that oligogenic inheritance may be linked to both male and female human infertility, and should therefore also be accurately measured when investigating male infertility.

In conclusion, in this article, we report the first evidence of oligogenic inheritance in altered spermatogenesis, leading to teratoasthenozoospermia. This mode of inheritance is crucial as oligogenic events could be behind the difficulties encountered by a significant proportion of infertile couples, for whom the current diagnosis is the somewhat unsatisfactory “unexplained” or “idiopathic” infertility. Our study was conducted in a context of teratozoospermia, a qualitative disorder of spermatogenesis. It paves the way for further studies on other male infertility disorders, including quantitative disorders, such as oligozoospermia or azoospermia for which the diagnostic yield remains very low. In particular, international consortia will benefit from this new understanding of the genetic factors influencing spermatogenesis by incorporating it into their scientific strategies. Current genetic tests to explore male infertility focus primarily on identifying low-frequency fully- penetrant monogenic defects, which are usually autosomal recessive and linked to the most severe cases of male infertility. However, investigation of oligogenic inheritance in the huge cohorts available should provide an estimate of the frequency of such events. These investigations could potentially identify new candidate genes involved in male infertility.

The discovery presented here is of major medical interest, and has implications for both clinical genetics and infertility management. First, the continuous and exponential characterization of new genes involved in infertility over the last decade offers the hope that, in the near future, an almost exhaustive list of genes and mutations involved in human infertility will be available. The identification of and screening for all known mutant alleles linked to male infertility at the heterozygous level should improve the diagnostic yield. Second, the discovery of multiple mutated genes will allow clinicians to provide more accurate genetic counselling to patients, and better guide them in their infertility journey. To improve patient management, future studies should look at potential correlations between patients’ mutational burden and their intracytoplasmic sperm injection (ICSI) success, pregnancies achieved, and live birth rates. It will also be essential to assess the impact of mutational load on parameters known to influence these outcomes.

## METHODS

### Animals

Generation of *Cfap43* and *Cfap44* KO mice is described in Coutton et al. (2018)^14^, generation of *Armc2* KO mice is described in Coutton et al. (2019)^26^. CRISPR/Cas9 gene editing was used to produce *Ccdc146* KO mice (ENSMUST00000115245). To maximize the chances of producing deleterious mutations, two gRNAs located in two distinct coding exons positioned at the beginning of the targeted gene were used. For each gene, the two gRNAs (5’-CCT ACA GTT AAC ATT CGG G-3’ and 5’-GGG AGT ACA ATA TTC AGT AC-3’) targeting exons 2 and 4, respectively, were inserted into two distinct plasmids also containing the Cas9 sequence. The Cas9 gene was driven by a CMV promoter and the gRNA and its RNA scaffold by a U6 promoter. Full plasmids (pSpCas9 BB-2A-GFP (PX458)) containing the specific sgRNA were ordered from Genescript (https://www.genscript.com/crispr-cas9-protein-crRNA.html). Both plasmids were co-injected into the zygotes’ pronuclei at a concentration of 2.5 ng/ml. It should be noted that the plasmids were injected as delivered by the supplier, thus avoiding the need to perform in vitro production and purification of Cas9 proteins and sgRNA. Several mutated animals were obtained with different insertions/deletions spanning a few nucleotides. All of the mutations obtained induced a translational frameshift expected to lead to complete absence of the protein or production of a truncated protein. The reproductive phenotype of two mutated lines was analyzed; homozygous males were infertile and displayed the same MMAF phenotype. For this study, a strain with a 4-bp deletion in exon 2 was used (c.164_167delTTCG).

For each KO strain, mice were maintained in the heterozygous state, and males and females were crossed to produce animals for subsequent generations. Heterozygous and homozygous animals were selected following PCR screening, the primers used for each strain are indicated in Supplementary Table 5. To produce multi-heterozygous animals, homozygous females were crossed with heterozygous males.

All animal procedures were conducted according to a protocol approved by the local Ethics Committee (ComEth Grenoble No. 318), by the French government (ministry agreement number #7128 UHTA-U1209-CA), and by the Direction Générale de la Santé (DGS) for the State of Geneva. Guide RNA, TracrRNA, ssDNA, and Cas9 were purchased from Integrated DNA Technologies. Pronuclear injection and embryo transfer were performed by the Transgenic Core facility at the Faculty of Medicine, University of Geneva. All genotypes were obtained by conventional interbreeding.

### Reproductive phenotyping

All adult male mice used were between 10 and 12 weeks old. After sacrifice by cervical dislocation, the testes were isolated and weighed, and sperm from caudae epididymides were allowed to swim for 10 min at 37 °C in 1 ml of M2 medium (Sigma-Aldrich, L’Isle d’Abeau, France). The sperm concentration was measured and adjusted before CASA analysis. An aliquot of sperm suspension was immediately placed in a 100-μm deep analysis chamber (Leja Products B.V., Nieuw-Vennep, the Netherlands), and sperm motility parameters were measured at 37 °C using a sperm analyzer (Hamilton Thorn Research, Beverley, MA, USA). The settings used for analysis were: acquisition rate: 60 Hz; number of frames: 45; minimum contrast: 50; minimum cell size: 5; low static-size gate: 0.3; high static-size gate: 1.95; low static-intensity gate: 0.5; high static-intensity gate: 1.3; minimum elongation gate: 0; maximum elongation gate: 87; magnification factor: 0.7. Motile sperm were defined by an average path velocity (VAP) > 1, and progressive sperm motility was defined by VAP > 30 and average path straightness > 70. A minimum of 200 motile spermatozoa were analyzed in each assay. Remaining sperm was rinsed with PBS-1X, centrifuged for 5 min at 500 g, spread on slides and allowed to dry at room temperature. Samples were then fixed in Ether/Ethanol 1:1 for Harris-Schorr staining (to assess overall morphology) or in 4% paraformaldehyde for DAPI staining (to assess nuclear morphology).

Morphology was assessed on a Nikon Eclipse 80i microscope equipped with a Nikon DS-Ri1 camera with NIS-ElementsD (version 3.1.) software. At least 200 spermatozoa were counted per slide at a magnification of ×1000.

### Analysis of nuclear morphology

Nuclear morphology was precisely evaluated by NMAS (version 1.19.2), according to the analysis method described in Skinner et al.^27^. The software processed images of DAPI-stained nuclei captured with a Zeiss Imager Z2 microscope, using a CoolCube 1 CCD camera, with a 100x/1.4 Zeiss objective and Neon software (MetaSystems, Altlussheim, Germany). Nucleus detection settings were Kuwahara kernel: 3, and flattening threshold: 100, for preprocessing; canny low threshold: 0.5, canny high threshold: 1.5, canny kernel radius: 3, canny kernel width: 16, gap closing radius: 5, to find objects; and min area: 1 000, max area: 10 000, min circ: 0.1, max circ: 0.9, for filtering. After acquisition of images of nuclei, landmarks were automatically identified using a modification of the Zahn-Roskies transform to generate an angle profile from the internal angles measured around the periphery of the nuclei. Angles were measured at every individual pixel around the original shape. This method combines data from every possible polygonal approximation into a single unified trace, from which landmark features can be detected (under-hook concavity, tail socket, caudal bulge and base, acrosomal curve, etc.). Angle profiles are presented as angle degrees according to the relative position of each pixel along the perimeter, and variability profiles use the interquartile range (IQR, difference between the third and first quartile) as a dispersion indicator to measure the variability of values obtained for each point. Sperm shape populations were then clustered using a hierarchical ward-distance method without reduction, based on angle profiles.

## STATISTICAL ANALYSIS

The statistics relating to nuclear morphology presented in Figure 8 and Supplementary Figure 4, were automatically calculated by the Nuclear Morphology Analysis Software. This analysis relied on a Mann-Whitney U test with Bonferroni multiple testing correction. p- values were considered significant when inferior to 0.05.

All other data were treated with R software (version 3.5.2). Histograms show mean ± standard deviation, and statistical significance of differences was assessed by applying an unpaired *t*- test. Statistical tests with two-tailed *p*-values ≤0.05 were considered significant.

## DATA AVAILABILITY

All data that support the findings of this study are available from the corresponding authors upon reasonable request.

## ACKNOWLEDGEMENTS

This work was supported in part by the French national research agency grants FLAGEL- OME: ANR-19-CE17-0014 and OLIGO-SPERM: ANR-21-CE17-00XX. We would like to sincerely thank Roxane DOMINGUEZ and Aurélien SIMON for their assistance with the software and IT solutions. We are also grateful to Charlotte GUYOT and Marlène GANDULA for technical assistance.

## AUTHOR CONTRIBUTIONS

G.M., C.C., P.F.R., and C.A. analyzed the data and wrote the manuscript; C.Ca., M.B., Z.- E.K., Y.N., and S.N. performed genetic work; G.M., C.L., E.L., M.D., G.C., C.V., J.M., J.-P.H., Y.N., S.N., and J.E. performed mouse work; G.M., C.C., P.F.R., and C.A. designed the study, supervised all laboratory work, had full access to all of the data from the study and take responsibility for the integrity of the data and its accuracy. All authors contributed to the report.

## COMPETING INTERESTS

The authors declare no competing financial interests.

## MATERIALS & CORRESPONDANCE

Requests should be addressed to G.M. and C.A.

## SUPPLEMENTARY INFORMATION

**Supplementary Figure 1.**
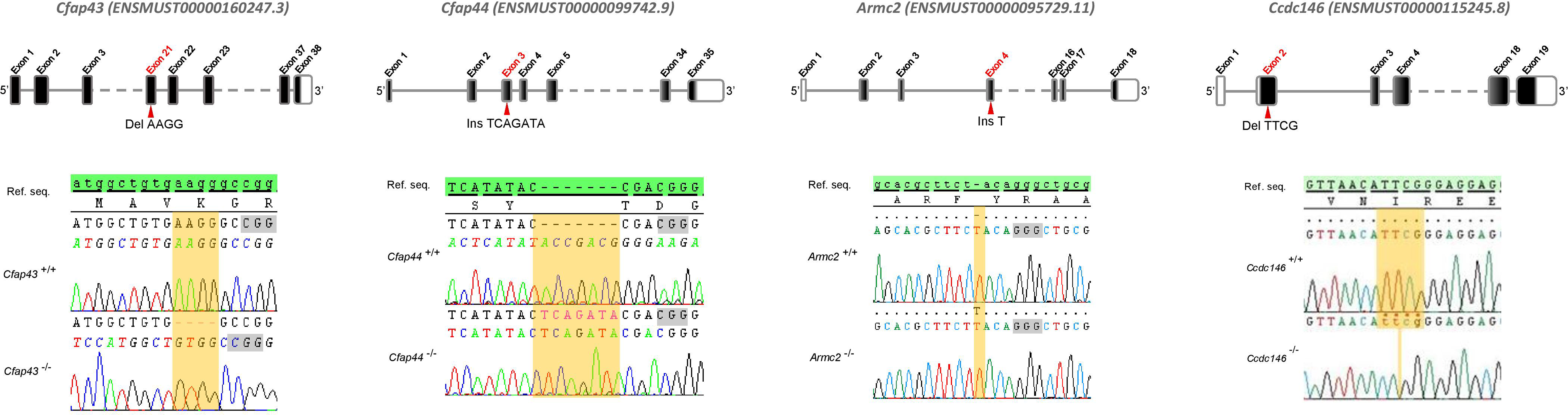
Location of mutations and electropherograms from Sanger sequencing of mutated forms of murine *Cfap43*, *Cfap44*, *Armc2*, and *Ccdc146* compared to their respective reference sequences. We confirmed a 4-bp deletion in *Cfap43* exon 21 (delAAGG), a 7-bp insertion in *Cfap44* exon 3 (InsTCAGATA), a 1-bp insertion (InsT) in *Armc2* exon 4, and a 4-bp deletion in *Ccdc146* exon 2 (delTTCG). Red arrows indicate the CRISPR/Cas9 targeting sequence, mutations are highlighted in yellow, and the gray boxes indicate the position of the protospacer-adjacent motif (PAM) sequences used during mutagenesis.

**Supplementary Figure 2.**
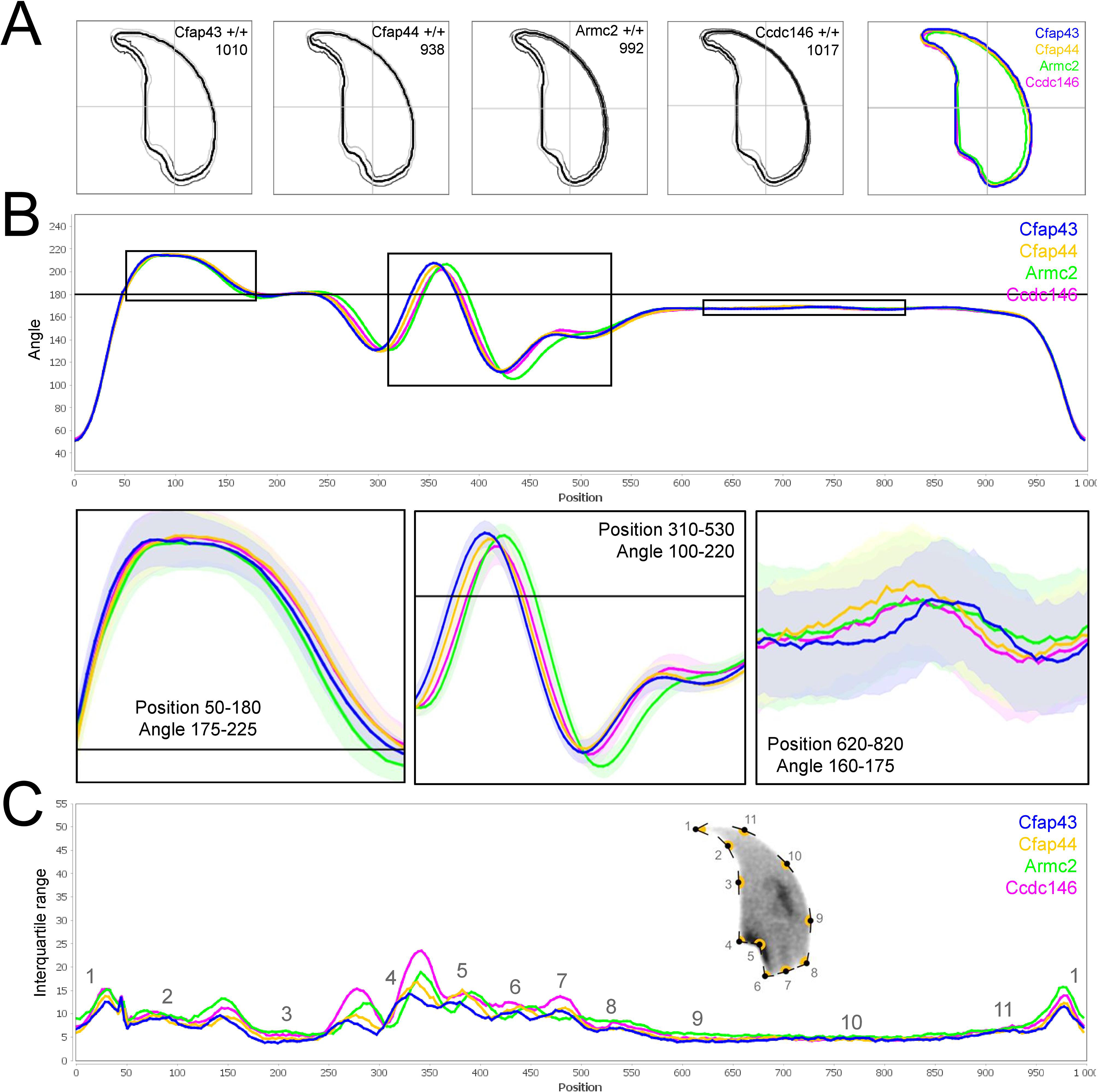
Comparison of fine nuclear morphology for wild-type animals produced from crosses of all strains studied. **A.** Consensus nuclear outlines for each strain alongside a merged consensus nucleus (blue=*Cfap43*, yellow=*Cfap44*, green=*Armc2* and pink=*Ccdc146*). The numbers assigned to each consensus outline correspond to the number of nuclei processed per condition. **B.** Angle profiles for each strain focusing (black boxes) on positions of specific interest. The x axis represents an index of the percentage of the total perimeter, as measured counterclockwise from the apex of the sperm hook. The y axis shows the interior angle measured across a sliding window centered on each index location **C.** Variability profiles for each strain. The x axis is the same as for the angle profile, and the y axis represents the Interquartile Range (IQR) (the difference between the 75th and 25th percentiles). Specific regions of the nuclei are mapped on the profile and the graphical representation (from ^27^), with: 1-tip; 2-under-hook concavity; 3-vertical; 4-ventral angle; 5-tail socket; 6-caudal bulge; 7-caudal base; 8-dorsal angle; 9-11-acrosomal curve.

**Supplementary Figure 3.**
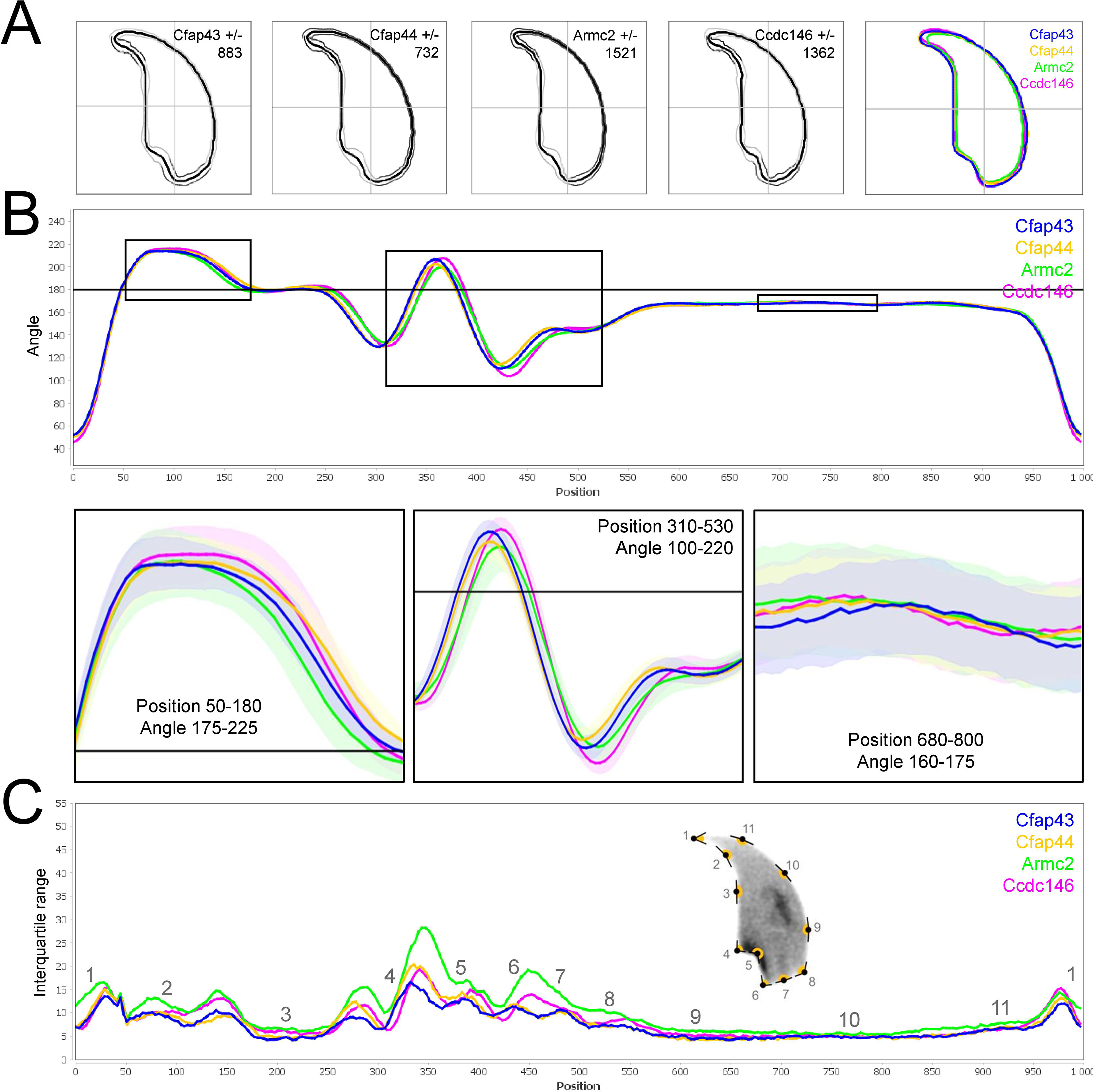
Comparison of fine nuclear morphology for heterozygous animals produced during crosses of all strains. **A.** Consensus nuclear outlines for each strain alongside a merged consensus nucleus (blue=*Cfap43*, yellow=*Cfap44*, green=*Armc2*, and pink=*Ccdc146*). The numbers assigned to each consensus outline correspond to the number of nuclei processed per condition. **B.** Angle profiles for each strain focusing (black boxes) on positions of specific interest. The x axis represents an index of the percentage of the total perimeter, as measured counterclockwise from the apex of the sperm hook. The y axis shows the interior angle measured across a sliding window centered on each index location **C.** Variability profiles for each strain. The x axis is the same as for the angle profile, and the y axis represents the Interquartile Range (IQR) (the difference between the 75^th^ and 25^th^ percentiles). Specific regions of the nuclei are mapped on the profile and the graphical representation (from ^27^) with: 1-tip; 2-under-hook concavity; 3-vertical; 4-ventral angle; 5-tail socket; 6-caudal bulge; 7-caudal base; 8-dorsal angle; 9-11-acrosomal curve.

**Supplementary Figure 4.**
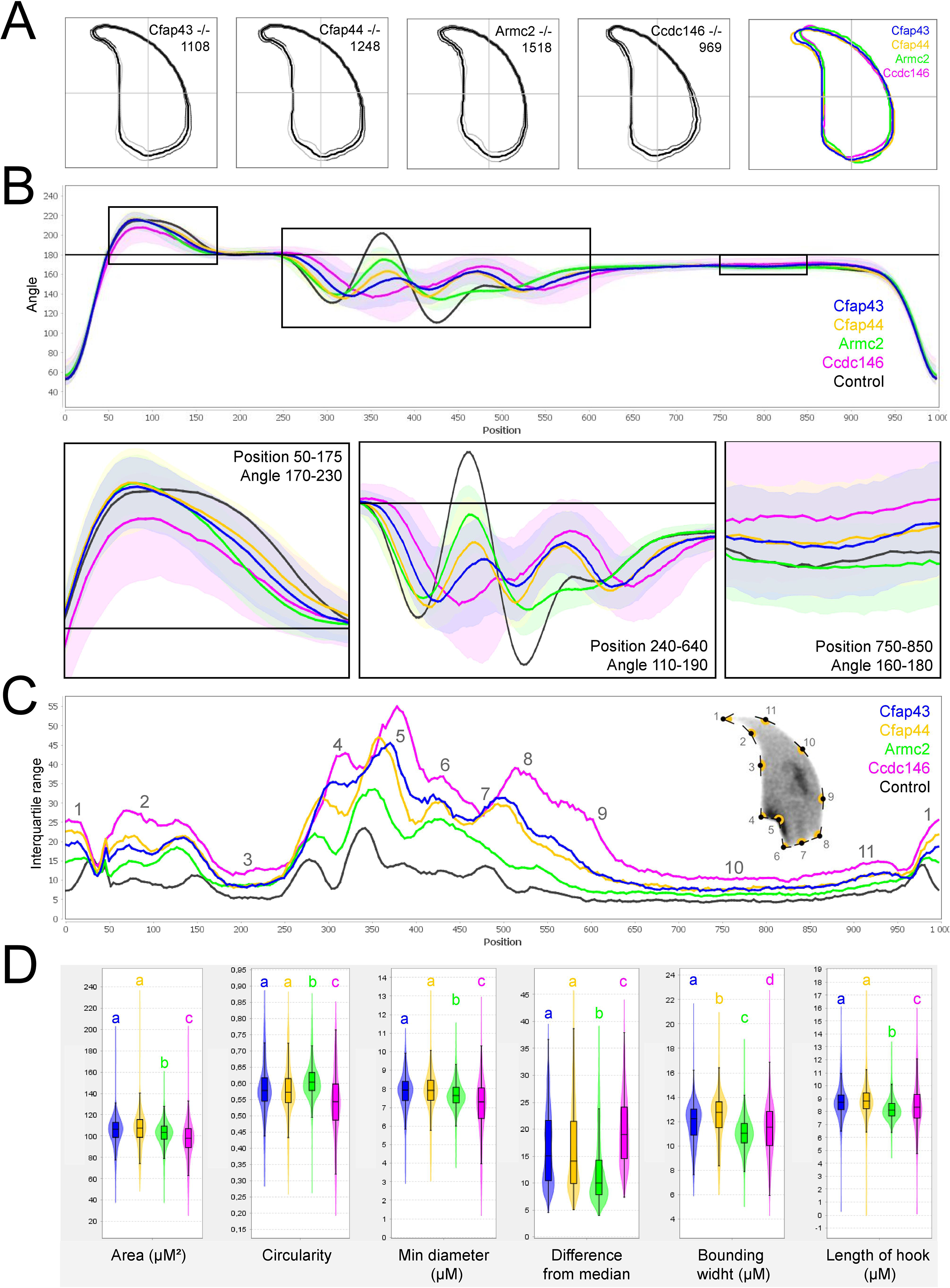
Comparison of fine nuclear morphology for sperm heads from KO and WT animals. **A.** Consensus nuclear outlines for each KO strain alongside a merged consensus nucleus, and (far right) superimposition of the outlines of the four strains (blue=*Cfap43*, yellow=*Cfap44*, green=*Armc2*, and pink=*Ccdc146*). The numbers assigned to each consensus outline corresponds to the number of nuclei processed per condition. **B.** Angle profiles for each strain alongside an angle profile for wild-type mice (black), focusing (black boxes) on positions of specific interest. The x axis represents an index of the percentage of the total perimeter, as measured counterclockwise from the apex of the sperm hook. The y axis corresponds to the interior angle measured across a sliding window centered on each index location **C.** Variability profiles for each strain. The x axis is the same as for the angle profile, and the y axis represents the Interquartile Range (IQR) value (the difference between the 75th and 25th percentiles). Specific regions of the nuclei are mapped on the profile and the graphical representation (from ^27^), with: 1-tip; 2-under-hook concavity; 3-vertical; 4-ventral angle; 5-tail socket; 6-caudal bulge; 7-caudal base; 8-dorsal angle; 9-11-acrosomal curve. **D.** Violin plots presenting mean data for representative nuclear parameters associated with each gene. Statistical significance of differences between populations was assessed by the NMAS software, applying a Mann-Whitney U test. Significantly different populations are identified by distinct letters.

**Supplementary Figure 5.**
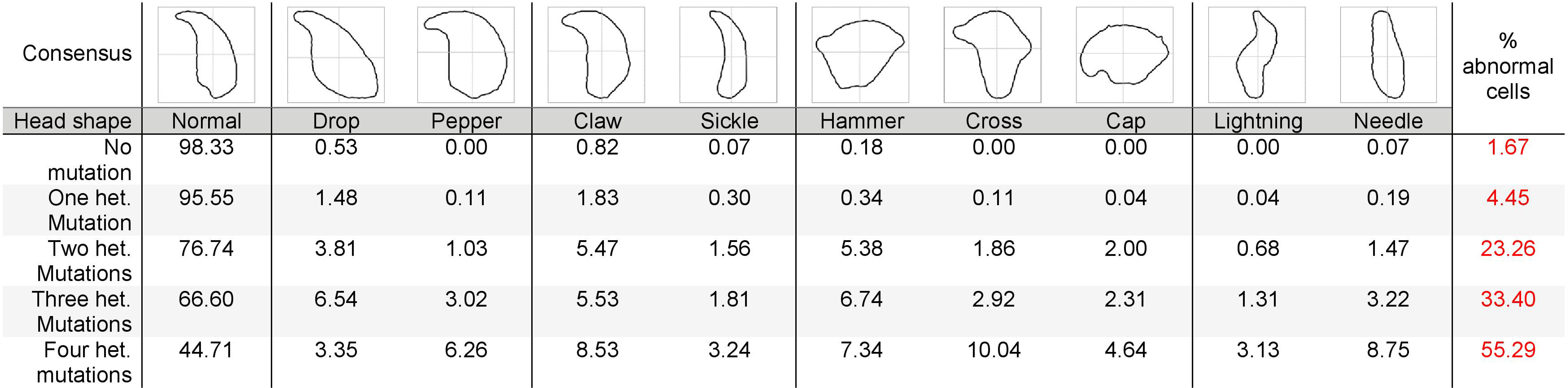
Consensus and percentage distribution of the distinct sperm populations classified based on nuclear morphology and according to the number of heterozygous mutations.

**Supplementary Table 1.** Statistical data linked to the Student t-tests performed in figure 5A-C. DoF = Degrees of Freedom; CI = Confidence Interval.

**Supplementary Table 2.** Statistical data linked to the Student t-tests performed in figure 5D-F. DoF = Degrees of Freedom; CI = Confidence Interval.

**Supplementary Table 3.** Statistical data linked to the Student t-tests performed in figure 6A-C. DoF = Degrees of Freedom; CI = Confidence Interval.

**Supplementary Table 4.** Statistical data linked to the Student t-tests performed in figure 6D-F. DoF = Degrees of Freedom; CI = Confidence Interval.

**Supplementary Table 5.** Statistical data linked to the Student t-tests performed in figure 7A. DoF = Degrees of Freedom; CI = Confidence Interval.

**Supplementary Table 6.** Statistical data linked to the Student t-tests performed in figure 7B. DoF = Degrees of Freedom; CI = Confidence Interval.

## Notes

### Competing Interest Statement

The authors have declared no competing interest.

